# Scaling up drug combination surface prediction

**DOI:** 10.1101/2024.12.24.630218

**Authors:** Riikka Huusari, Tianduanyi Wang, Sandor Szedmak, Diogo Dias, Tero Aittokallio, Juho Rousu

**Affiliations:** Department of Computer Science, Aalto University, P.O. Box 11000 (Otakaari 1B), FI-00076, Espoo, Finland; Institute for Molecular Medicine Finland (FIMM), HiLIFE, University of Helsinki, Helsinki, Finland; Hematology Research Unit Helsinki, University of Helsinki and Helsinki University Hospital, Helsinki, Finland; Translational Immunology Research Program, University of Helsinki, Helsinki, Finland; Institute for Cancer Research, Department of Cancer Genetics, Oslo University Hospital, N-0310, Oslo, Norway; Oslo Centre for Biostatistics and Epidemiology (OCBE), Faculty of Medicine, University of Oslo, N-0317, Oslo, Norway

**Keywords:** drug combination prediction, drug interaction surfaces, kernel methods, structured output prediction

## Abstract

Drug combinations are required to treat advanced cancers and other complex diseases. Compared to monotherapy, combination treatments can enhance efficacy and reduce toxicity by lowering the doses of single drugs – and there especially synergistic combinations are of interest. Since drug combination screening experiments are costly and time consuming, reliable machine learning models are needed for prioritizing potential combinations for further studies. Most of the current machine learning models are based on scalar-valued approaches, which predict individual response values or synergy scores for drug combinations. We take a functional output prediction approach, in which full, continuous dose-response combination surfaces are predicted for each drug combination on the cell lines. We investigate the predictive power of the recently proposed comboKR method, which is based on a powerful input-output kernel regression technique and functional modelling of the response surface. In this work, we develop a scaled-up formulation of the comboKR, that also implements improved modeling choices: 1) we incorporate new modeling choices for the output drug combination response surfaces to the comboKR framework, and 2) propose a projected gradient descent method to solve the challenging pre-image problem that traditionally is solved with simple candidate set approaches. We provide thorough experimental analysis of comboKR 2.0 with three real-word datasets within various challenging experimental settings, including cases where drugs or cell lines have not been encountered in the training data. Our comparison with synergy score prediction methods further highlights the relevance of dose-response prediction approaches, instead of relying on simple scoring methods.

## 1 Introduction

Drug combination therapies are increasingly needed for treating cancer and other complex diseases [1–3]. A number of studies have used the various computational tools of bioinformatics for better understanding of mechanism of diseases and treatment of different diseases. Recently published in-silico studies showed that bioinformatic play as vital role in drug development and and better understanding of the molecular basis of diseases [4–8]. However, systematic discovery of synergistic drug combinations remains challenging, due to limited capacity and cost of experimental drug screening. Even with high-throughput screening technologies [9–12], exhaustive testing of pairwise combinations leads to increasingly large combinatorial search space of drug combinations; for instance, considering one hundred drugs at five concentrations in ten cell lines would result in more than one million laboratory tests. In this regard, machine learning approaches can speed-up and rationalize the search process by pinpointing the most promising search directions [13, 14].

While most of the machine learning research focuses on prediction of drug combination synergy scores or other single-valued prediction problems [14–18], a few recent approaches consider scalar-valued machine learning approaches for predicting directly the drug combination responses at selected concentrations [19–23]. These methods are more applicable in practice and to various combination prediction tasks beyond synergy, as they are not dependent on any specific synergy metric. Indeed, the different synergy models that score the divergence between the expected and measured combination response – such as the highest single agent (HSA) model [24], Bliss independence model [25] or Loewe additivity model [26] – do not always agree in terms of synergy definition, partly due to large differences in drug concentrations and maximum response values across experimental studies [27]. The approaches that predict individual response values have severe limitations. For instance, predictions for a given drug combination might not be consistent, nor is there any guarantee that they would follow a smooth response surface. Indeed, individual response values can be seen as samples of an underlying target function: a non-linear parametric surface describing the drug interactions. By directly predicting such functional form, the models provide better interpolation and extrapolation along the surfaces, and more consistently capture and quantify potential synergistic patterns.

To this end, [28] and [29] proposed models that predict full drug interaction surfaces, instead of discrete values sampled from them. Of these structured prediction approaches, PIICM introduced in [28] is based on Gaussian process regression, working similarly to a matrix completion approach in filling-in missing drug surface entries from the batch of experiments. ComboKR [29] belongs to the family of functional output regression methods that predict the drug interaction surfaces from primary features, such as gene expression of cell lines or structural fingerprints of drugs.

The comboKR approach is based on a powerful input-output kernel regression framework [30–32], with a novel normalisation scheme for the drug interaction surfaces [29]. In this work, we propose an improved formulation for the comboKR framework that both scales up to larger datasets, and obtains a predictive performance superior to the original approach; we call this scalable variant comboKR 2.0. In addition to the improved scalability, we propose two additional improvements to the model: 1) we introduce the usage of difference to the neutral interaction surfaces as the intermediate target for the learner, and 2) we solve the arising pre-image problem, that is often solved with simple candidate set optimisation, with a more advanced projected gradient descent algorithm. We perform comprehensive computational experiments to validate and benchmark our approach, showing results in three different drug combination datasets and three predictive scenarios of varying difficulties. We also perform ablation studies, showcasing the relevance of our proposed model improvements, as well as perform comparison against two popular deep learning -based synergy score prediction approaches.

## 2 Methods

### 2.1 ComboKR

ComboKR, first introduced in [29], uses generalised kernel dependency estimation (KDE) [32] or input output kernel regression (IOKR) [30, 31] framework to predict full, continuous drug interaction response surfaces *y* ∈ 𝒴 from the input data *x* ∈ 𝒳. The idea of the framework, in a nutshell, is to cast the difficult difficult structured learning problem *f* : 𝒳 → 𝒴 into a simpler learning problem. This is done by mapping the structured data in 𝒴 to a reproducing kernel Hilbert space (RKHS) ℋ_𝒴_ associated with a kernel chosen for the output data, *k*_*y*_ : 𝒴 *×* 𝒴 → **R**. This gives rise to a vector-valued^1^ learning problem to learn *g* : 𝒳 → ℋ_𝒴_ . After learning *g*, a (usually ill-posed) pre-image problem has to be solved to map the predictions back to 𝒴.

ComboKR uses kernel ridge regression (KRR) in learning the function *g*, with which a closed-form solution is obtained and directly inserted to the final pre-image problem. With this, and assuming a normalised output kernel *k*_*y*_ (i.e. *k*_*y*_(*y, y*) = 1 ∀*y* ∈ 𝒴), the final optimisation problem to obtain predictions can be written as

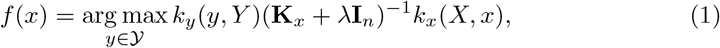

in which **K**_*x*_ is the *n×n* kernel matrix for *n* input training samples; **I**_*n*_ is identity matrix of same size; *λ* is the KRR regularisation parameter; and the shorthand *k*_*y*_(*y, Y*) refers to the vector [*k*_*y*_(*y, y*_1_), …, *k*_*y*_(*y, y*_*n*_)] with 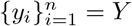 the outputs of the training set; *k*_*x*_(*X, x*) is defined analogously. It is good to note, that here the kernel trick is used for both input and output data. For the output data, a crucial part of the comboKR approach was to propose a novel drug interaction surface normalisation scheme within the output kernel *k*_*y*_, that captures the relevant parts for the similarity comparisons with it. It was demonstrated that this approach was much more effective than the use of a more traditional kernel computation.

The search space 𝒴 in pre-image problem is vast, and the problem is ill-posed [33, 34]. ComboKR solved the problem with candidate set optimisation: instead of considering the full 𝒴, the search is restricted to finding maximum value over a smaller candidate set *C*_*y*_. Every element in the set is tried out, and the one giving maximum value to 1 is chosen as the final prediction.

As is usual for kernel methods, the solution depends on inverting a *n × n* matrix, **K**_*x*_ + **I**, with *n* the number of training samples. In general the input data space would consist of triplets (*x*_*c*_, *x*_*d*1_, *x*_*d*2_) ∈ 𝒳 = 𝒳_*c*_ × 𝒳_*d*_ × 𝒳_*d*_, with cell line features and the features from the two drugs in the combination. In comboKR, the model was built for one cell line at a time, and input data consisted of tuples (*x*_*d*1_, *x*_*d*2_) ∈ 𝒳_*d*_ × 𝒳_*d*_. Now **K**_*x*_ = **R**[**K**_*d*_ ⊗**K**_*d*_]**R**^⊤^, with 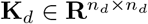 begin kernel matrix of *n*_*d*_ drug features, and 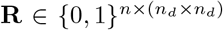 selects the *n* training samples from the full Kronecker product matrix. For the full problem including also cell lines, **K**_*x*_ = **R**^*′*^[**K**_*c*_ ⊗ **K**_*d*_ ⊗ **K**_*d*_]**R**^*′*⊤^ with 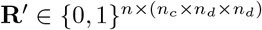, which is not practical to compute for most datasets.

### 2.2 Scaling up ComboKR

At the first glance, scaling up the inverse computation might seem trivial - in the end the kernel matrix on the full data is based on simple Kronecker product **K**_*c*_ ⊗ **K**_*d*_ ⊗ **K**_*d*_. However the training set is rarely complete, and **K**_*x*_ does not include a full set of cell-drug-drug combinations. In practice, for training one gets a sampled version of this matrix, which does not enjoy the efficient inverse formula taking advantage of the Kronecker product structure (eq. (512) in [35]).

A solution is to use efficient iterative solvers for the KRR problem. Such solver has been proposed in [36, 37], with practical implementation in [38]. This iterative generalised vec-trick solver is able to take advantage of the sampled Kronecker product structure of the input kernel matrix, and solve a linear equation of the form

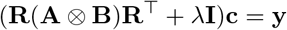

by efficient computation of the iterative update **a** ← **R**(**A** ⊗ **B**)**R**^⊤^**c** with sparse **R**. However, the method is only proposed for products consisting of two matrices in the product. Thus with the drug response prediction application, a part of the three-part Kronecker product has to be explicitly computed before applying the solver; **K**_*c*_ ⊗ **K**_*d*_ or **K**_*d*_ ⊗ **K**_*d*_, depending on if number of cell lines or drugs in the data is smaller.

Using the iterative solver to obtain the KRR solution for function *g* : 𝒳 → ℋ_𝒴_ means, that it is no longer possible to take advantage of the kernel trick in the output space: now for a data sample *x* the prediction step is done by solving the pre-image problem

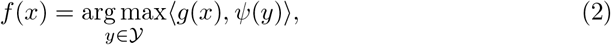

where *Ψ* denotes the feature map associated with the normalised output kernel *k*_*y*_.For some kernels, such as the simple linear kernel, *Ψ*(*y*) is easily computable, but for example for the RBF kernel used in comboKR, the feature space is infinite-dimensional.

In order to be able to compute (2), we need to consider approximation of this output space, 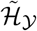, and learn with KRR instead 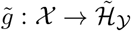

To this end, we consider a Nyström approximated feature map of the output surface kernel *k*_*y*_. Generating a diverse set of landmark surfaces, 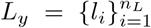, that cover the space well and give a good quality approximation, we can compute 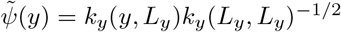, giving the final pre-image problem as

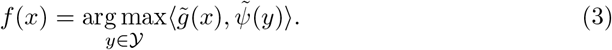

Most of the drug combinations in the datasets exhibit behaviour close to additive effects, i.e. not antagonistic or synergistic combinations. We can write a generic surface function as *S*(*c, d*_1_, *d*_2_, *κ*) → ℝ, depending on the cell line and the two drugs (implicitly encoding the monotherapy behaviour) and some interaction parametrisation *κ*. Intuitively, when looking at the resulting surfaces, any small changes in the interaction pattern can be drowned into the more prominent overall patterns of the surfaces. Since the additive part can be directly deduced from the monotherapy responses, there is no need to use the capacity of the learner to learn it, but it can rather be focused on only the interaction pattern. Thus, in order to be able to better capture the tails of the uneven target distribution (i.e. the antagonistic or synergistic surfaces), we propose to write *S*(*c, d*_1_, *d*_2_, *κ*) = *S*(*c, d*_1_, *d*_2_, 0) + Δ, and focus predicting only Δ, i.e. the difference of the true and the neutral interaction surfaces. The final prediction is obtained by adding the neutral surface *S*_0_(*c, d*_1_, *d*_2_, 0) back to the predicted Δ.

This is similar modeling choice as in [28], where the surfaces were also partitioned to a non-interaction term and an interaction term that captured the possible synergistic or antagonistic effect.

#### Algorithm 1

Pre-image problem solver to obtain prediction for input *x* associated with cell line *c* and drugs *d*_1_ and *d*_2_.

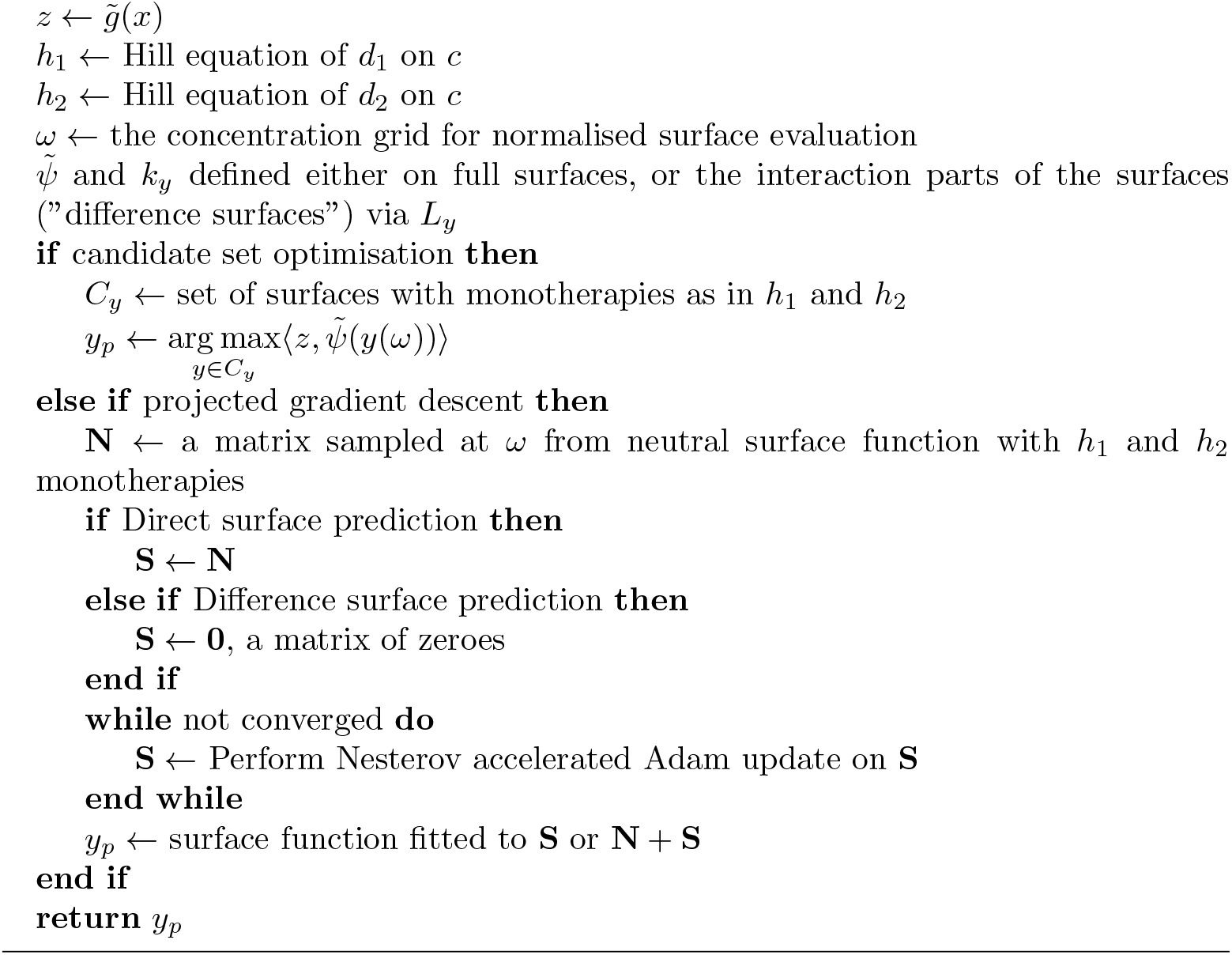

When solving the pre-image problem, the simplest approach is again to use the candidate set, and try out different interaction parametrisations *κ* in *S*(*c, d*_1_, *d*_2_, *κ*). In addition to this, we consider projected gradient descent algorithm. More concretely, we use Nesterov accelerated Adam optimisation strategy, and project the result back to valid surfaces by fitting the surface model to the end-result of the gradient descent. Due to the rarity of antagonistic or synergistic interactions, a good first guess for the prediction is a surface without any interaction effects. Thus, the gradient descent uses either *S*(*c, d*_1_, *d*_2_, 0) or zero-valued function as the initial starting point, depending if the learning is done directly on full surfaces, or only on interaction part (Δ) of it. The algorithms are displayed in Algorithm 1.

The comboKR 2.0 pipeline is illustrated in Figure 1, where also the proposed improvements compared to the original comboKR approach are highlighted.

**Fig. 1.**
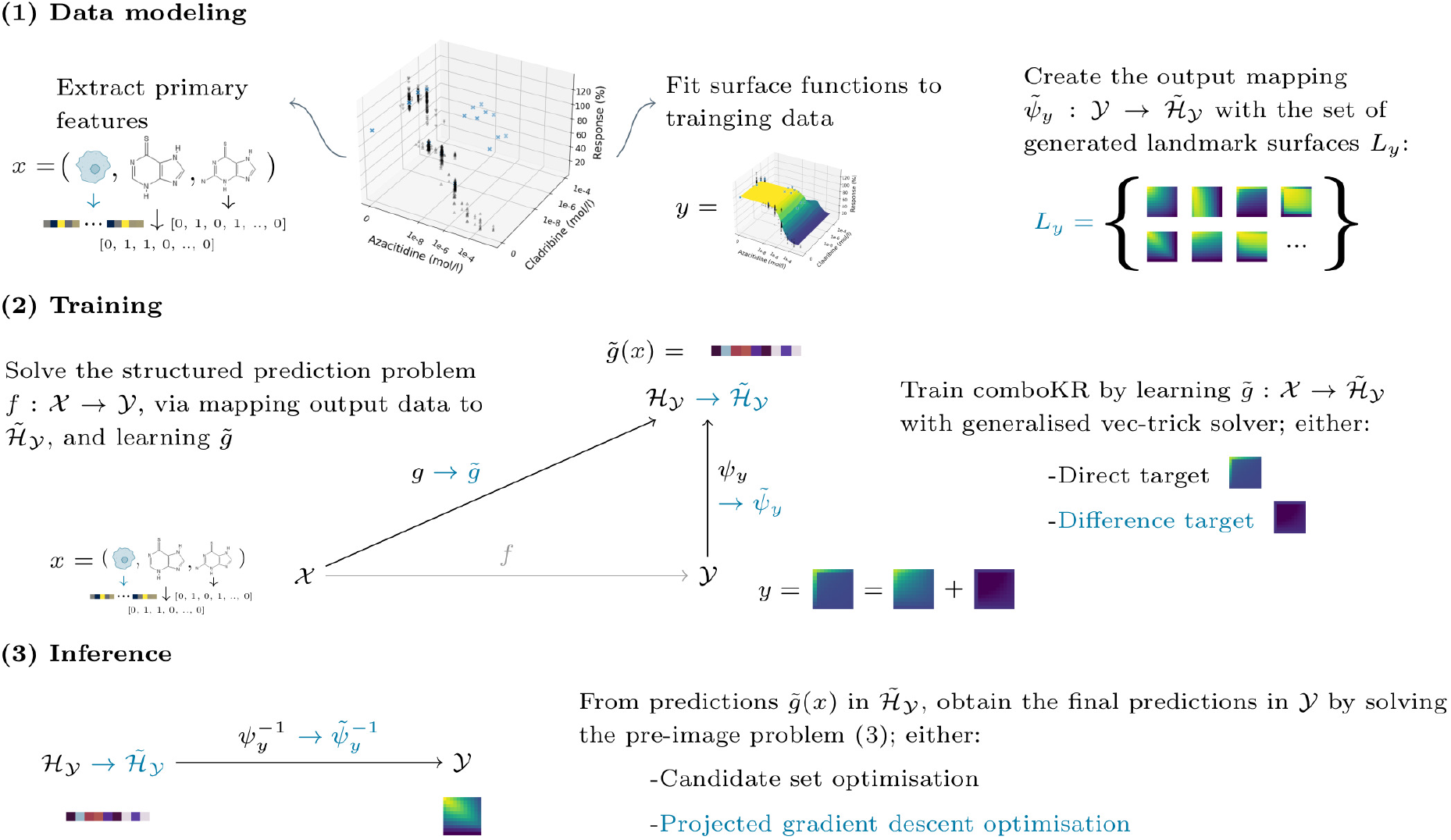
Illustration of the comboKR 2.0 approach to solve the function-valued drug surface prediction problem. First the surface model is fitted to the training data, and landmarks are generated in order to form the required approximation of the output space. The comboKR model is trained by using the generalised vec trick solver. Predicting requires solving the challenging pre-image problem. The improvements compared to the previous comboKR approach are highlighted with teal-coloured text.

## 3 Datasets

We consider three datasets – Jaaks et al [11], NCI-ALMANAC [10] and O’Neil et al [39] – collecting drug combination dose-response measurements on various cancer cell lines. The details of the various datasets are described below, while a summary of relevant statistics of them is displayed in Tables 1 and 2.

**Table 1.** The number of cell lines and drugs in the datasets considered in this work. The table reports the amount and sizes of the drug combination matrices measured in the data – the percentage of surfaces is calculated over all possible combinations (in Jaaks data a surface count does not differentiate between anchor and library drugs).

| Dataset | #Cell lines | #Drugs | #Surfaces | Measurements |
| --- | --- | --- | --- | --- |
| Jaaks et al | 125 | 64 | 68 982 ($\approx 27.4\%$) | $7 \times 2$; one or two |
| NCI-ALMANAC | 60 | 104 | 311 527 ($\approx 96.9\%$) | $3 \times 3$; one set |
| O’Neil et al | 39 | 38 | 22 527 ($\approx 82.2\%$) | $4 \times 4$; four sets |

**Table 2.**
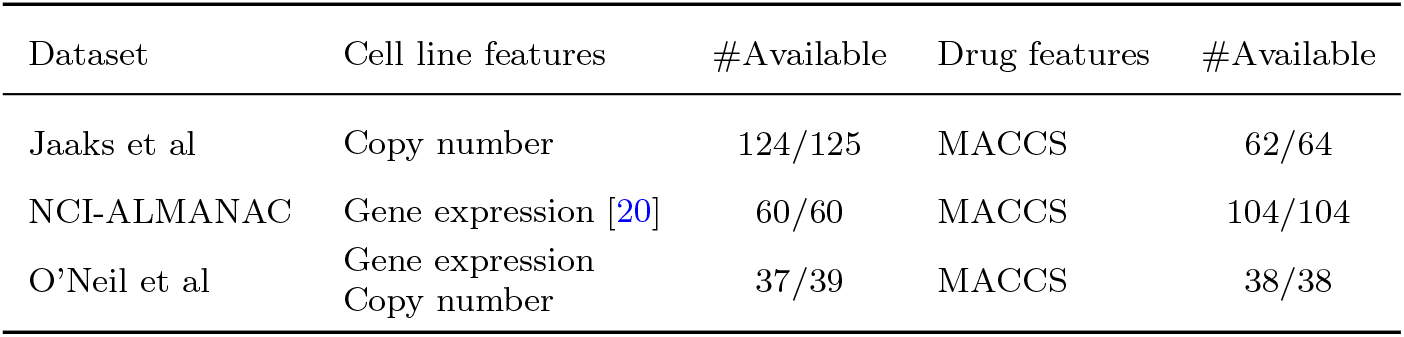
The primary features used with the various datasets, with number of cell lines and drugs for which they are available.

| Dataset | Cell line features | #Available | Drug features | #Available |
| --- | --- | --- | --- | --- |
| Jaaks et al | Copy number | 124/125 | MACCS | 62/64 |
| NCI-ALMANAC | Gene expression [20] | 60/60 | MACCS | 104/104 |
| O’Neil et al | Gene expression<br>Copy number | 37/39 | MACCS | 38/38 |

### 3.1 Jaaks et al

The dataset consists of testing 64 drugs on 125 cell lines, with the cell lines being from breast, colon or pancreatic tumors. More specifically, for the three tumor types, 56, 26 and 25 unique drugs were considered, respectively. The response data consists of 7 *×* 2 drug combination measurements done with an anchored approach: the anchor drug with pre-selected low and high doses was tested against a range of seven increasing concentrations of the second compound. The matrices with the same anchor and library drugs were considered as additional monotherapy data, and were not included in the surfaces.

For primary features, we collected the corresponding 125 DepMap ids for the cell lines, and downloaded copy number data for 124 of the cell lines. Similarly for drugs, we searched for PubChem ids, with which we collected the SMILES representation for 62 out of 64 drugs, and computed MACCS structural features.

### 3.2 NCI-ALMANAC

The NCI-ALMANAC dataset is based on testing 104 drugs in 60 cell lines from 9 tissue origins. The data consists of 3 × 3 drug combination measurements on the cell lines; in total there are 311 527 drug-drug-cell combinations. In addition to the 3 × 3 combination responses, the data also collects monotherapy responses for the corresponding concentrations, providing hence 4 × 4 response matrices when they are included.

As primary features for the cell lines, we considered the gene expression data from [20]. As primary features for the drugs, we use the MACCS fingerprints calculated from the isometric SMILES representation.

### 3.3 O’Neil

The dataset is built on testing 38 drugs on 39 cell lines representing multiple cancer types. The data consists of 4 × 4 drug combination measurements on the cell lines; in total there are 22 527 drug-drug-cell combinations.^2^ Median response values over the replicate measurements are considered as the ground truth responses for the experiments.

As the primary features for the drugs, the MACCS fingerprints were calculated based on the isometric SMILES representation.

As primary features for the cell lines, we consider the gene expression profile data and copy number from DepMap [40]. These data were not obtained for two of the cell lines.

### 3.4 Surface model fitted to the data

As in [29], in this work, we consider the BRAID (Bivariate Response to Additive Interacting Doses) drug interaction model [41] that builds on the Hill equation [42, 43] and is motivated by the Loewe additivity principle. Indeed, most of the parameters of the model are the Hill equation parameters for the monotherapy behaviour of the two drugs, with just two interaction parameters.

As comboKR heavily depends on good estimates of the underlying drug response surfaces, Figure 2 displays the optimal fit of the BRAID surfaces achieved to all the available data. The responses range from 0 to either 1 or 100, representing the percent growth of the cells. For better comparison across the different data sources, for this figure the O’Neil dataset responses have been transformed to [0, 100] from the original [0, 1].

**Fig. 2.**
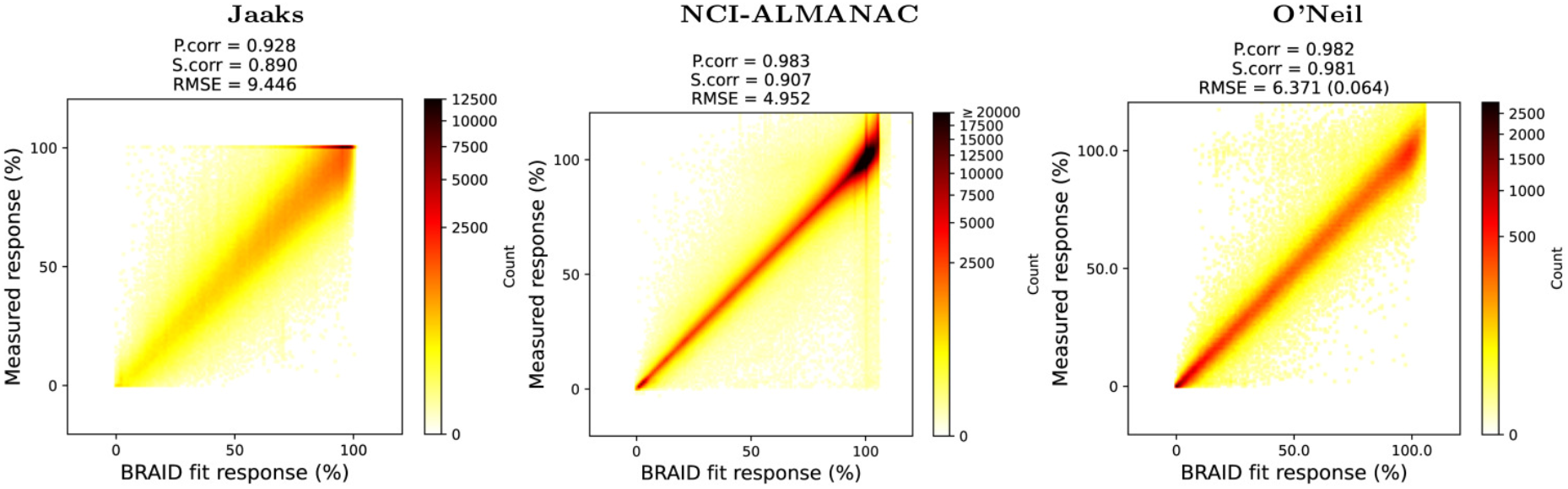
Density plots of the fitted BRAID functions sampled at the dose-concentrations of the measurements, compared to the measured responses. With Jaaks data, if a drug combination occurs in two ways in a cell line, only one of them has been included. For O’Neil data, the median of replicate measurements are used as groundtruth, and the response values have been transformed from the [0, 1] range to [0, 100] range for better comparison between the datasets; the RMSE value in parenthesis corresponds to the original smaller response value range [0, 1].

Of the datasets considered, O’Neil achieves the best fit in all metrics, and overall has most even distribution of the response values over the full range. In contrast, both NCI-ALMANAC and Jaaks data contain more responses where the drugs do not inhibit the cell growth.

## 4 Computational experiments

### 4.1 Predictive scenarios

We consider three predictive scenarios, here listed in order of increasing difficulty:

- **New combo:** a prediction is done on a new triplet (*c, d*_1_, *d*_2_), where the cell line and both drugs have been seen in the training set as parts of other triplets, but the triplet itself not.
- **New drug:** the triplet (*c, d*_1_, *d*_2_) contains either a novel pair (*c, d*_1_) or (*c, d*_2_): one of the drugs in the triplet has not been seen in the training set associated with that cell line.
- **New cell line:** at the prediction time, the triplet (*c, d*_1_, *d*_2_) queried contains a cell

line not seen in the training set. However, the drug combinations in test set can appear in the training data.

### 4.2 Competing methods

We compare our proposed scaled-up comboKR to the one originally proposed in [29], and denote the proposed scalable variant as comboKR 2.0^3^.

Modeling and the drug combination responses as a structured prediction problem is still an emerging topic. Apart from the previous comboKR approach [29], to our knowledge only PIICM [28] has been proposed in this context. While giving promising results in a simple predictive scenario, it is however not suitable in more challenging ones, as it assumes at the test stage that all cell lines and drugs are present in training data. In addition, due to suboptimal scalability to larger datasets, it is not possible to offer meaningful comparisons in the present experiments.

LTR is a polynomial regression method based on latent tensor reconstruction [44], that has been successfully applied to drug combination response prediction [23] as “comboLTR”. In its original form, it is applicable to settings with one data view, or with multiple views they can be concatenated (early fusion approach). However, this “one-view” approach is not scalable to large datasets. Thus, a more scalable variant has been proposed in [45], that allows to handle magnitudes larger data sets than the original version. We note that even with same parameters, the polynomials learned by the one-view and multi-view comboLTR models are not the same, as the interactions are modeled differently. Due to the scalability issues of the one-view LTR, we use it here only with the O’Neil dataset. Multi-view LTR is considered with all the datasets.

As a simple baseline approach, we compare to BRAID surfaces without any synergy or antagonism. These surfaces are assigned without any learning, and they are based on the same monotherapy Hill equation parameters that the comboKR/comboKR 2.0 uses.

While our setting is designed for predicting response surfaces, we additionally compare our method with two synergy score prediction methods: DeepSynergy [17] and MatchMaker [18]. Both of these models are popular baseline methods for the drug combination synergy prediction tasks based on feedforward neural network architectures; the DeepSynergy model was one of the first deep learning approaches proposed in the synergy score prediction literature.

In our comparison with the synergy score prediction models, we predict the response surfaces with comboKR and comboLTR as usual, and subsequently compute the synergy scores as a postprocessing step. DeepSynergy and MatchMaker are directly trained with synergy scores; we consider both Bliss and Loewe synergies separately. In order to have only one synergy score per surface, we take the mean of the synergy scores of the available combinations dose-responses (e.g. for O’Neil dataset over the 16 combo measurements of the surface, similarly for the other data sets; see Table 1).

### 4.3 Experimental details

ComboKR relies on kernels on both inputs and outputs. As mentioned, for outputs we consider RBF kernel on the normalised surfaces sampled on a grid. For the input features, either Tanimoto or RBF kernel is calculated – Tanimoto is used for the fingerprint features, RBF kernel for others. In all RBF kernels, the kernel parameter *σ* is set as mean of pairwise distances in the data.

For non-kernel based approaches (i.e. ComboLTR), the empirical features based on the input kernels are considered. Additionally, as a scalar-valued prediction approach, ComboLTR requires concentration values. These are given as one-hot encoded vectors.

Cross-validation is performed to select the regularisation parameters for the models. For the various methods, the parameters searched over are as follows:

- comboKR 2.0: we consider early stopping of the iterative generalised vec-trick solver as the regularisation; the number of iterations are cross-validated over [3, 5, 10]. Number of landmark BRAID surfaces in Nyström approximated output feature map can also been as a regulariser in all the models. It’s decided by the number of eigenvalues greater than or equal to 0.1 in the kernel formed from generated landmark surfaces. This results in only 27 landmark surfaces in Nyström approximation. Unless otherwise stated, we consider difference surface modeling and projected gradient descent as the pre-image solver.
- Original comboKR: regularisation parameter *λ* in IOKR: [1e-2, 5e-2, 1e-1, 5e-1]
- ComboLTR: in both variants order of the polynomial is chosen from [3, 5], and rank from [10, 20].
- We perform the cross-validation for DeepSynergy and MatchMaker over learning rate, and cross-validate over learning rates [0.01, 0.001, 0.0001, 0.00001].

## 5 Results

### 5.1 Overall comparison

We present the overall Pearson correlation results of the compared approaches in Figure 3. The comboKR 2.0 outperforms the previous comboKR approach; only with O’Neil data and on the easiest new combo scenario, the original comboKR perform bettern than 2.0. The major improvement from the original is seen especially clearly in the new drug experimental scenario, where the comboKR 2.0 outperforms the baseline, while original version obtains worse results than the baseline. Overall, comboKR outperforms the comboLTR in all predictive scenarios. As can be expected, the new cell line scenario is very challenging, and the difference of comboKR 2.0 to the baseline is very small.

**Fig. 3.**
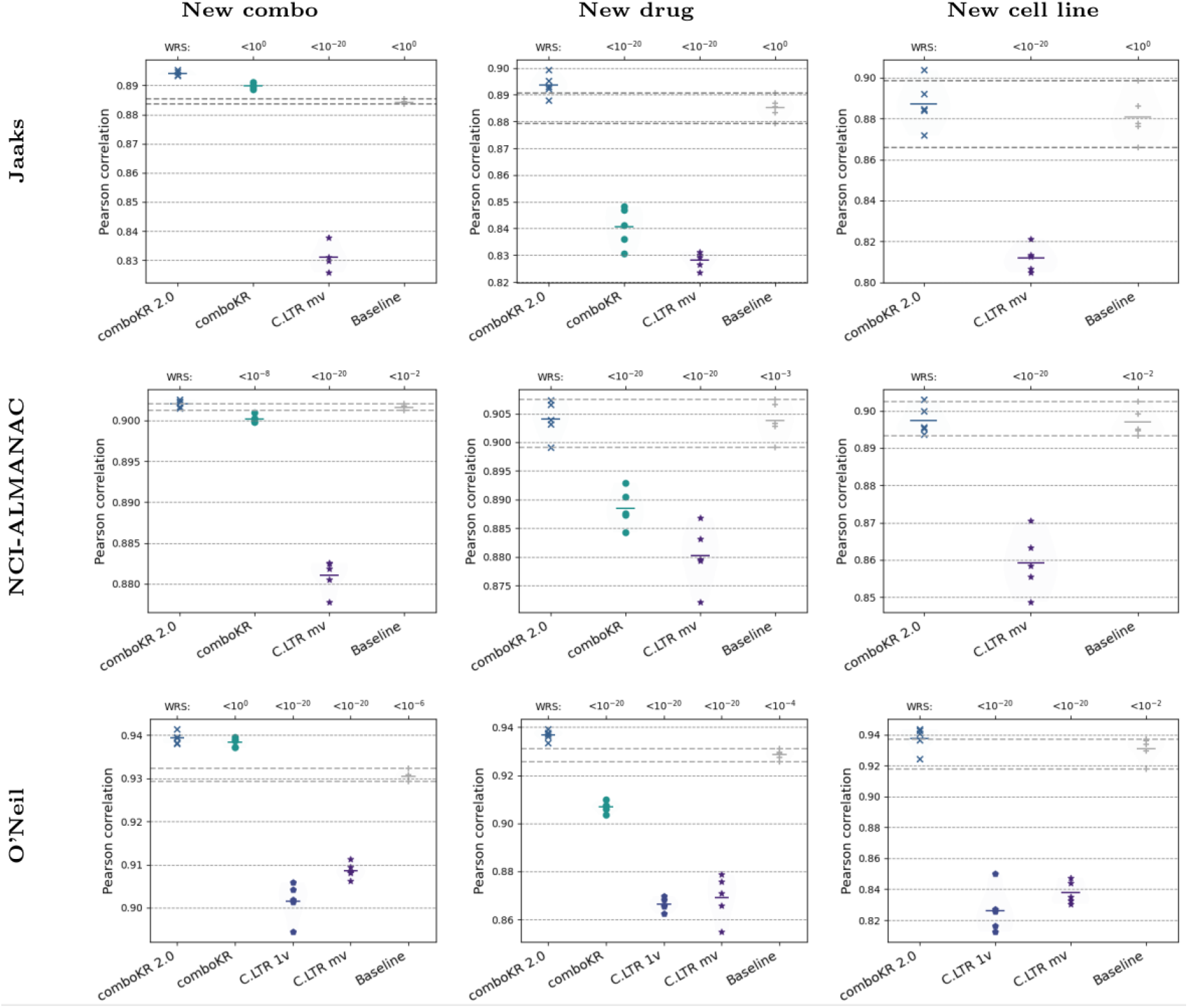
Pearson correlation results of the various methods on the three datasets, between the measured and predicted drug combination responses. The original ComboKR model is trained cell-by-cell manner, while the other methods are trained over multiple cell lines: thus, comboKR is not applicable in the new cell line scenario. The horizontal dashed lines highlight the range of baseline results of predicting neutral interaction surface without learning via the BRAID function (last entries in the plots). On top of the plots the p-value results of Wilcoxon rank-sum test (“WRS”) are shown (with Bonferroni correction), compared to our comboKR 2.0.

In order to perform statistical testing on the results, we computed for each surface prediction its root mean squared error (RMSE) to the groundtruth surface measurements. With these, we performed Wilcoxon rank-sums test, as well as Kolmogorov-Smirnov test, comparing the other methods to our proposed comboKR 2.0. We display the Bonferroni-corrected p-values as part of Figure 3.

### 5.2 Comparison with synergy score prediction methods

As drug combination synergy prediction methods outnumber the drug combination response prediction methods, we here present comparison between the two types of methods on the O’Neil dataset. Here, we consider two types of synergy models: Loewe and Bliss. We present the Pearson correlation results between the predicted synergy score and the synergy score computed from the ground truth data in Figure 4. There, we also show the results of the Wilcoxon rank-sum test with Bonferroni correctio, where for all the methods, we compared the distributions of the predicted scores to the ground truth scores, taking the median over the five splits. We note, that for the dose-response prediction models (comboKR, comboLTR), the synergy score prediction is obtained via a post-processing step, where the scores are calculated based on the predicted responses.

**Fig. 4.**
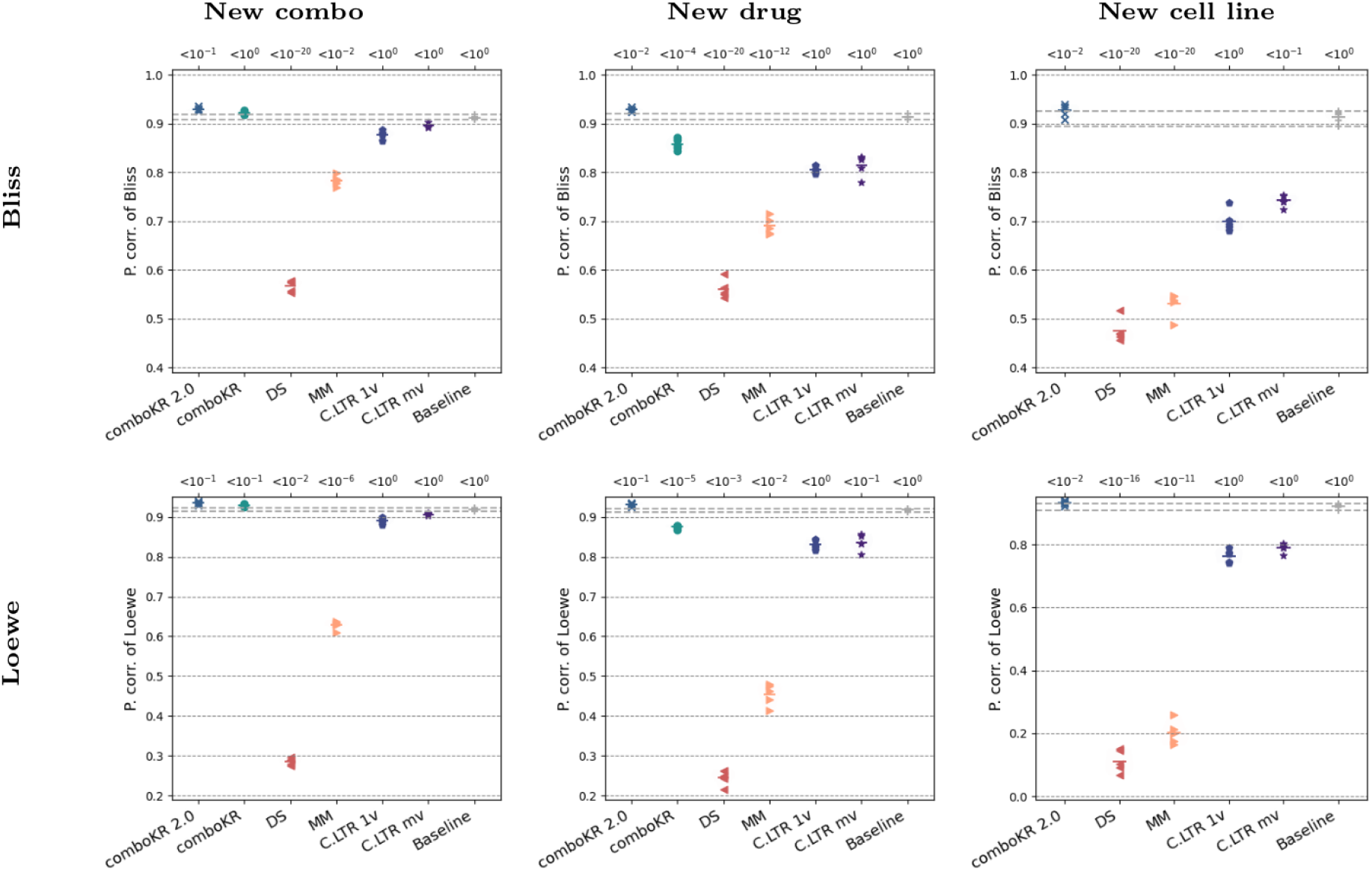
Pearson correlations of the synergy score prediction results on O’Neil data, compared to the scores computed from the ground truth measurements. On top of the plots them median (over the five splits) p-value results of Bonferroni-corrected Wilcoxon rank-sum test are shown, for when the synergy score predictions are compared to the ground truth scores.

The results higlight the relevance of dose-response prediction methods. While the synergy score prediction method MatchMaker is competitive in the new combo experimental setting with Bliss synergy score, the same method performs much worse when Loewe is considered. Indeed, it seems that Loewe synergy scores are much more difficult than Bliss for these methods. Deep learning methods often thrive with large data sets sizes; it could be expected that the performance of DeepSynergy and Match-Maker would improve with larger data set sizes, or with more detailed cross-validation strategies than our comparison considered.

### 5.3 Cell line feature importance

In addition to the importance of even including the cell line features, we briefly investigate the performance with different cell line features with O’Neil dataset. To this end, we consider using three different primary features for the cell lines: A) the gene expression and copy number features used in other experiments; B) one-hot encoding of the cell line id; and C) one hot encoding of the cell line id, concatenated with one-hot encoding of the cancer type id.

The results are displayed in Figure 5. In the easier new combo training scenario all methods benefit from using id features; comboLTR seems to additionally benefit from using the cell type id features. For comboKR 2.0, the performance of using cell type and cell line ids is here equal to that using more complicated primary features. The new drug scenario starts to show how the complicated primary features can be a better choice than the id features, while the difference between the features in comboLTR also decreases.

**Fig. 5.**
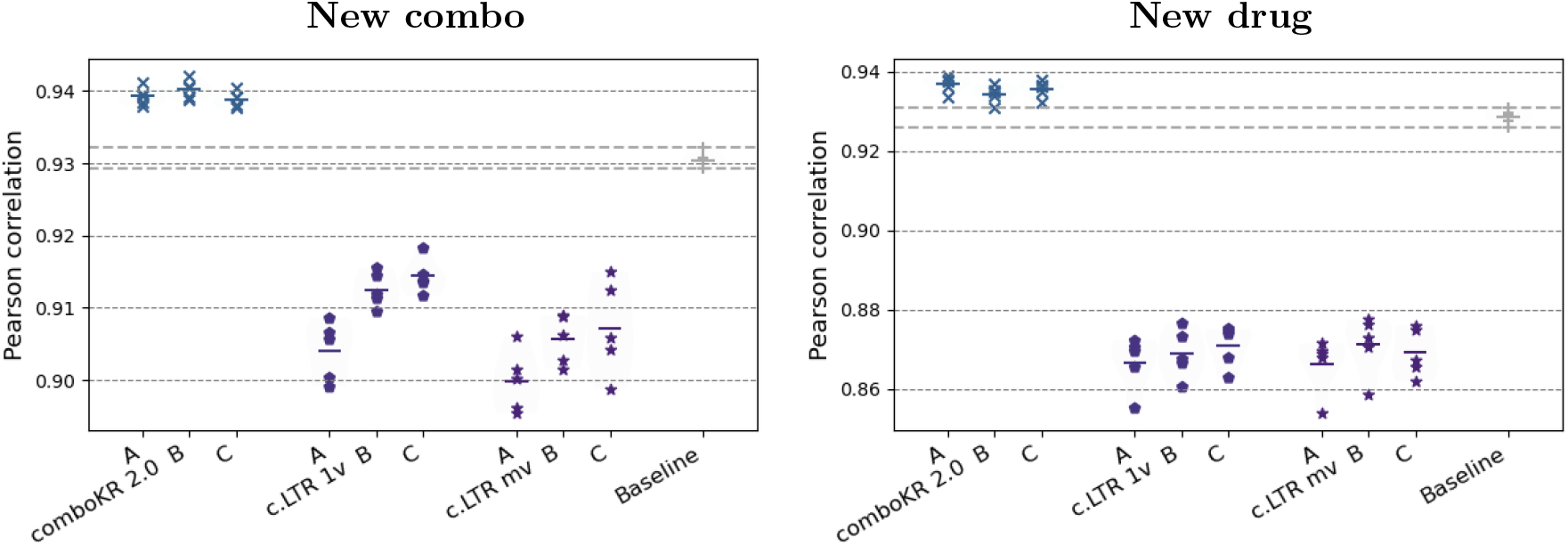
Pearson correlations of the response predictions with various cell line features on O’Neil data: A: primary features (gene expression and copy number features) B: one-hot encoded cell line id C: one-hot encoded cell id with one-hot encoded cancer type id.

### 5.4 Ablation study

The scaled-up version of comboKR often outperforms the original implementation that considers each cell line a separate experiment. In order to investigate how much of the gain/loss is due to larger training set sizes, the scaled-up algorithms were run also in cell-by-cell manner and compared to the full runs.

The Pearson correlation of the full and cell-by-cell comboKR 2.0 approach are shown in Table 3. It seems clear that the improved modeling choices improved the comboKR 2.0 approach, independently from scaling up for larger data set sizes. Furthermore, the results indicate that it is possible to obtain good results with comboKR 2.0 even when only data available is from the investigated cell line – an important improvement for practical applications.

**Table 3.** Pearson correlation results (the mean ± standard deviation) on the comboKR 2.0 when trained and tested on full training set, compared to the cell-by-cell setting, and original comboKR.

| Dataset |  | comboKR 2.0 | c.KR 2.0 cbc | c.KR orig. |
| --- | --- | --- | --- | --- |
| Jaaks | New combo | $0.894 \pm 0.001$ | $0.895 \pm 0.000$ | $0.890 \pm 0.001$ |
| | New drug | $0.893 \pm 0.004$ | $0.889 \pm 0.004$ | $0.839 \pm 0.007$ |
| NCI-ALMANAC | New combo | $0.902 \pm 0.000$ | $0.902 \pm 0.000$ | $0.900 \pm 0.000$ |
| | New drug | $0.904 \pm 0.003$ | $0.904 \pm 0.003$ | $0.889 \pm 0.003$ |
| O’Neil | New combo | $0.939 \pm 0.001$ | $0.940 \pm 0.001$ | $0.938 \pm 0.001$ |
| | New drug | $0.937 \pm 0.002$ | $0.934 \pm 0.002$ | $0.907 \pm 0.002$ |

We further investigated substituting the more advanced projected gradient descent solver on difference surface modeling with the simpler modeling possibilities and algorithms to solve the pre-image problem. The different parametrisations with or without difference surface target, and with the different pre-image approaches. We denote these as follows:

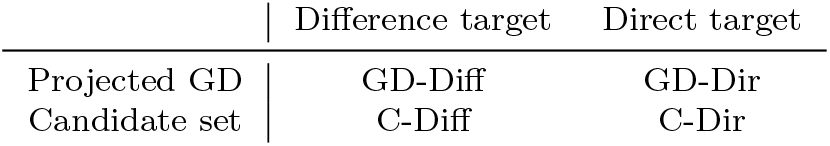

The results are displayed in Figure 6. It is clear that considering difference surfaces within the comboKR model provides consistently better results than using the original surface as the target. Interestingly, using projected gradient descent as the pre-image solver improves the performance further with difference surfaces, but significantly worsens the results with original surface modeling.

**Fig. 6.**
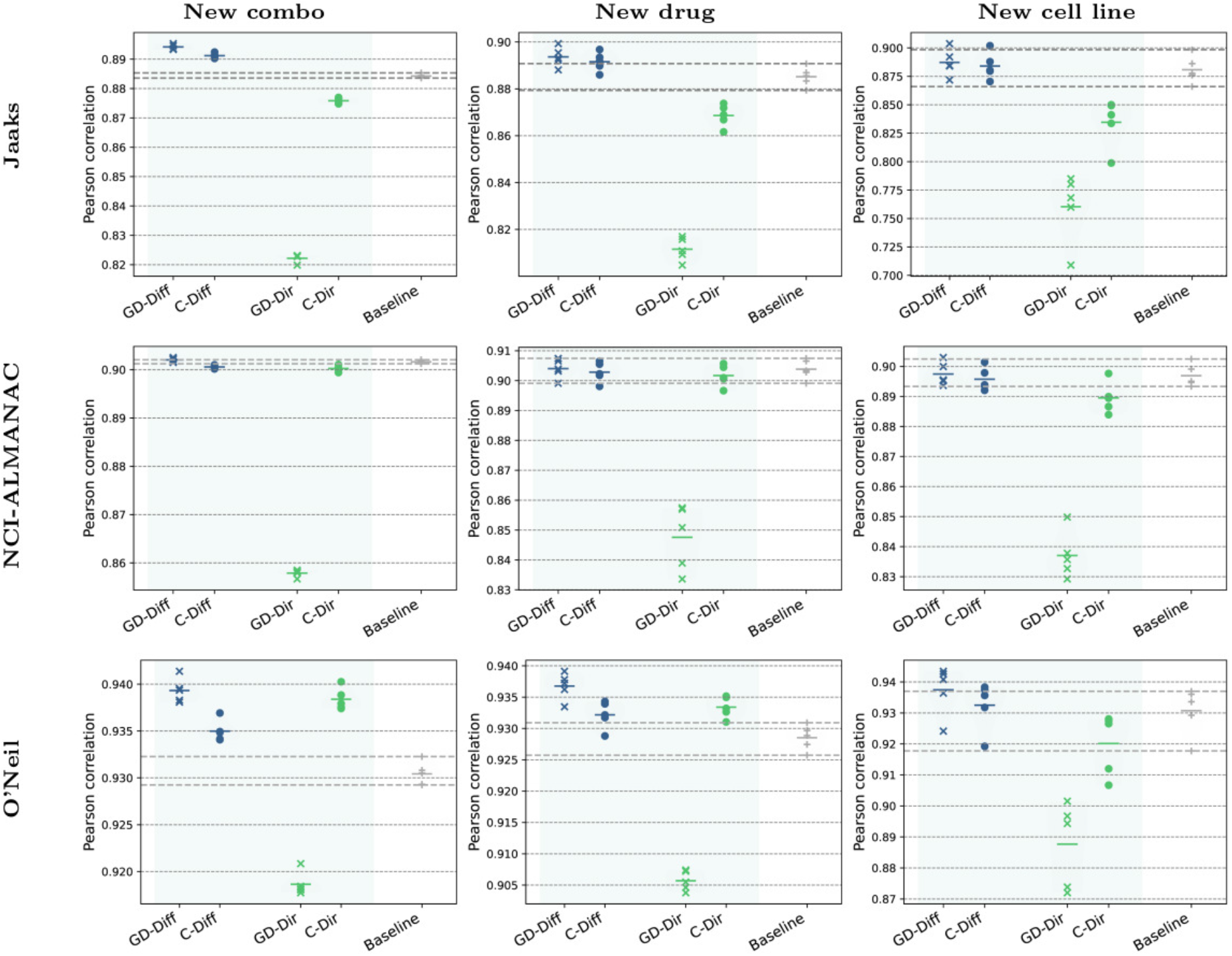
Pearson correlation results of the comboKR 2.0 variants on the three datasets, between the measured and predicted drug combination responses.

## 6 Discussion

In this work, we have implemented multiple improvements to the recently proposed comboKR method [29], a novel way to predict dose responses as full drug combination surfaces. The improvements include scalability to larger datasets with data from multiple cell lines, as well as improvements in the model configuration and approaches for solving the arising pre-image problem. Due to the combinatorial nature of the drug combination response datasets, the scalability of the predictive methods will be very important in the future: as one or two new drugs or cell lines are added to experiments, the amount of response-surfaces grows in combinatorial fashion.

Our work demonstrates the effectiveness of the improved comboKR 2.0 against the original version, as well as against relevant baseline methods, in many relevant experimental scenarios of varying difficulty. The results on three real-world datasets highlight especially the suitability of the variant with difference surface modeling in various experimental designs, coupled with projected gradient descent pre-image optimisation. Our method offers competitive performance compared to the competing comboLTR model as well as the original comboKR, and our ablation study highlights the relevancy of the proposed modeling choices. Indeed, with difference surface modeling, the method is able to take advantage of the projected gradient descent pre-image optimisation method, unlike in the original direct surface prediction setting, where candidate set optimisation far outperforms gradient descent. While comboKR 2.0 excels in the difficult new drug scenario, improved cell line feature modeling might still lead to better performance in this most challenging scenario of predicting responses in previously unseen cell lines.

As a functional output prediction approach – compared to approaches that predict individual drug combination dose-responses – our approach offers predictions that are consistent across the full dose-response surfaces. Moreover, from the predicted surface it is easy to obtain predictions to any dose-concentrations of interest, or compute any type of synergy measure of interest. As illustrated by our experimental comparison to synergy score prediction methods, our comboKR 2.0 is very competitive in synergy score prediction, even if the objective is to predict responses, not the synergy scores. While the proposed comboKR 2.0 method seems to outperform the other methods, it is important to note that neutral interaction surfaces from Baseline (i.e. additive BRAID surface set without learning), are often extremely competitive: indeed, strong synergism or antagonism is a relatively rare phenomenon [3, 11, 46, 47]. This is especially well seen with the NCI-ALMANAC dataset, which is much more complete than the other two datasets (see Table 1). This large-scale data is therefore expected to contain much more simple additive surfaces, and we found with this dataset most challenging to outperform the additive baseline.

As the case with the initial comboKR, also comboKR 2.0 heavily relies on good surface model to be used in both training and testing time. While the BRAID model has provided promising results, in the future other surface models, such as MuSyC [48] or generalised Hill-type response surfaces [49], could be investigated in our framework to allow for more diverse surface representations.

## Data availability

The original datasets are available for downloading as follows:

- Jaaks [11]:https://figshare.com/articles/dataset/Original_screen_All_tissues_raw_data_csv_zip/19141916?file=34010816
- NCI-ALMANAC [10]:https://wiki.nci.nih.gov/display/NCIDTPdata/NCI-ALMANAC
- O’Neil [39]:https://aacrjournals.org/mct/article/15/6/1155/92159/An-Unbiased-Oncology-Compound-Screen-to-Identify

The BRAID functions fitted to the data, as well as the Hill equations and the cross-validation splits of the different experiments are available at zenodo: https://zenodo.org/records/14509938; DOI: 10.5281/zenodo.14509938.

## Acknowledgements

The authors acknowledge the computational resources provided by the Aalto Science-IT project.

## Footnotes

1 or function-valued for kernels associated with infinite-dimensional RKHS

2 In total there are 22 727 combinations in the data, however 200 of them are not measured at a 4 × 4 grid, so we excluded those in favour of easy evaluation on regular grid size.

3 Code available at https://github.com/aalto-ics-kepaco/comboKR2.0

